# Long Read Sequencing reveals transgene concatemerization and backbone integration following AAV-driven electroporation of CRISPR RNP complexes in mouse zygotes

**DOI:** 10.1101/2024.02.18.580906

**Authors:** Muhammad W. Luqman, Piroon Jenjaroenpun, Jessica Spathos, Nikhil Shingte, Mitchell Cummins, Pattaraporn Nimsamer, Lars M. Ittner, Thidathip Wongsurawat, Fabien Delerue

**Affiliations:** Dementia Research Centre, Macquarie Medical School, Faculty of Medicine, Health and Human Sciences, Macquarie University, Sydney, Australia; Khyber Medical University, Institute of Medical Sciences, Kohat, Pakistan; Division of Medical Bioinformatics, Research Department, Faculty of Medicine Siriraj Hospital, Mahidol University, Bangkok, 10700, Thailand; FOXG1 Research Foundation, Sands Point, New York, USA; School of Biotechnology and Biomolecular Sciences, the University of New South Wales, Sydney, Australia

## Abstract

Over the last decade CRISPR gene editing has been successfully used to streamline the generation of animal models for biomedical purposes. However, one limitation to its use is the potential occurrence of on-target mutations that may be detrimental or otherwise unintended. These bystander mutations are often undetected using conventional genotyping methods.

The use of Adeno-Associated Viruses (AAVs) to bring donor templates in zygotes is currently being deployed by transgenic cores around the world to generate knock-ins with large transgenes. Thanks to a high level of efficiency and the relative ease to establish this technique, it recently became a method of choice for transgenic laboratories. However, a thorough analysis of the editing outcomes following this method is yet to be developed.

To this end, we generated three different types of integration using AAVs in two different murine genes (i.e., *Ace2* and *Foxg1*) and employed Oxford Nanopore Technologies long read sequencing to analyze the outcomes. Using a workflow that includes Cas9 enrichment and adaptive sampling, we showed that unintended on-target mutations (sometimes reported using other workflows) can occur when using AAVs. This work highlights the importance of in-depth validation of the mutant lines generated and informs the uptake of this new method.

## Introduction

Genetically modified (GM) animals, particularly mice, are powerful models to understand the mechanisms underlying physiological processes and human disorders. They are also invaluable tools to develop and test novel treatment strategies.

Transgenic laboratories around the world generate these models for biomedical research purposes, using either microinjection techniques (Delerue and Ittner 2017), or more recently electroporation of fertilized zygotes (Qin et al. 2015; Kaneko and Mashimo 2015). Electroporation is less challenging than microinjection and allows for high-throughput transformation of zygotes, whereas microinjection requires manipulation of zygotes one at a time. Moreover, survival and development rates are comparatively higher because electroporation is less invasive (Kaneko and Mashimo 2015). As such, electroporation of zygotes is widely used to generate knockouts (KO) and small nucleotides changes, such as point mutations or base pair exchanges. We, and others, generated such mouse models using electroporation of one-cell embryos (Klugmann et al. 2022; Morey et al. 2022).

However, electroporation remains largely inefficient at driving the targeted integration of large transgenes to generate knock-in (KI) mouse lines, presumably because the zona pellucida (thick glycoprotein membrane protecting the embryos at the preimplantation stages) prevents the shuttling of these large transgenes inside the embryos. To the best of our knowledge, there has been no report to date of a successful transformation of double stranded DNA (e.g., plasmid or transgene) using electroporation of such zygotes, and the largest insertion reported so far by electroporation of single-stranded oligo is 1kb in length (Miyasaka et al. 2018).

Therefore, a new method based on infection of zygotes with Adeno-Associated Viruses (AAVs) to bring the donor template inside the zygotes, coupled with electroporation of CRISPR ribonucleoprotein (RNP) complexes to induce Homology Directed Repair (HDR) has recently been developed (Chen et al. 2019). Such method, coined “CRISPR-READI” shows high level of efficiency (up to 100% in our hands) and is relatively easy to implement in transgenic laboratories. Romeo *et al*. showed that AAVs could diffuse through the zona pellucida (Romeo et al. 2020), while the Rivera-Perez lab demonstrated that the transfection efficiency varies depending on the serotype used, serotype 6 having one of the highest levels of transduction in mouse zygotes (Yoon et al. 2018).

Over the last decade CRISPR gene editing has been extensively used to generate GM mice, however, one limitation to its use is the potential occurrence of on-target mutations that are detrimental or otherwise unintended. These bystander mutations are typically undetected using conventional genotyping (i.e., PCR) and routine (i.e., Sanger) sequencing (Simkin et al. 2022; Sailer et al. 2021). Recently, Oxford Nanopore Technologies (ONT^©^) Long Read Sequencing (LRS) has been used in mice to identify the insertion site of randomly integrated transgenes (Bryant et al. 2023), and to confirm integration following recombinase-mediated cassette exchange (RMCE) (Low et al. 2022). However, to the best of our knowledge, LRS has not yet been used to perform quality control (QC) following AAV-driven gene editing in zygotes. To this end, we performed CRISPR-READI to generate three different types of integration in two different murine genes (i.e., *Ace2* and *Foxg1*). We then analyzed these knock-ins using the Oxford Nanopore Technologies (ONT^©^) MinION Mk1c and/or GridION, following a Cas9-based enrichment method that we previously applied to cells (Wongsurawat et al. 2020). Using this method, we identified instances of concatemerization with partial backbone integration (particularly Inverted Terminal Repeat sequences - ITR) in two out of five (40%) mouse lines generated.

## Results

### Generation of KI mouse lines

We performed AAV-driven gene editing on two different murine genes: the angiotensin I converting enzyme 2 (i.e., *Ace2*) and the forkhead box G1 (i.e., *Foxg1*) to insert three different transgenes.

First, we targeted the start codon of the murine *Ace2* gene and inserted the human Ace2 (hAce2) coding sequence upstream of a polyadenylation signal (SV40pA) (Fig. 1a).

**Figure 1.**
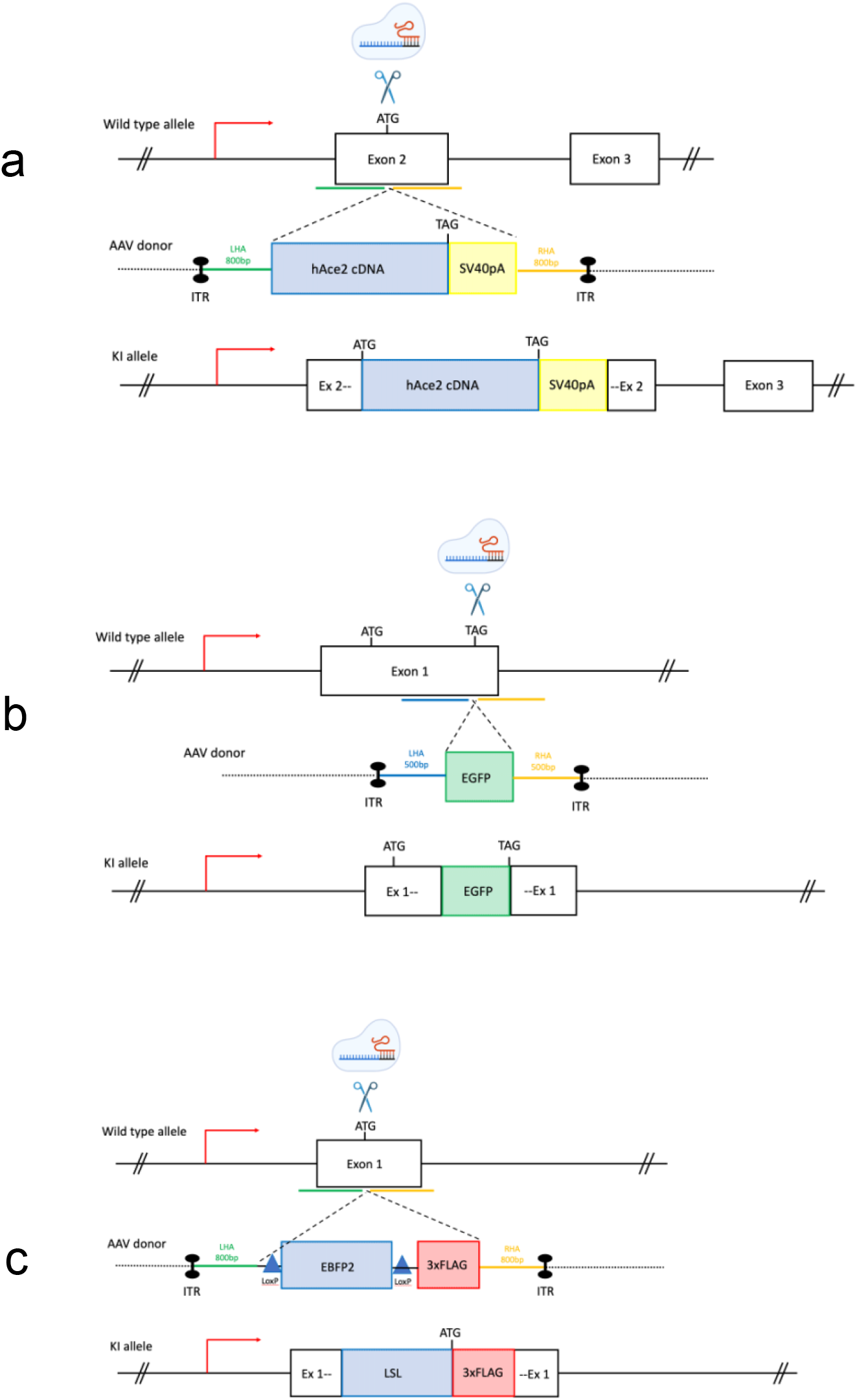
Generation of KI mouse lines by AAV-driven gene editing. The gene editing strategy for each knock-in (KI) line is illustrated. (a) To generate the humanized Ace2 (hAce2) mice, editing reagents targeting the start codon of the murine *Ace2* gene were electroporated in fertilized zygotes, following AAV-driven delivery of the donor template containing the hAce2 coding DNA sequence upstream of an PolyA. (b) CRISPR sgRNAs were directed to the strop codon of the mouse *FOXG1* gene to fuse an EGFP sequence at the c-terminus. (c) The start codon of the murine *FOXG1* gene was targeted with sgRNAs to insert a stop cassette and a triple FLAG tag. LHA = left homology arm, RHA = right homology arm, ITR = inverted terminal repeat sequence, hAce2 = human Ace2 coding sequence, SV40pA = simian virus 40 polyadenylation signal, EGFP = Enhanced Green Fluorescence Protein sequence, EBFP2 = Enhanced Blue Fluorescent Protein sequence, 3xFLAG = triple FLAG tag, LSL = Lox-Stop-Lox.

We next targeted the stop codon of the endogenous *Foxg1* gene to insert an Enhanced Green Fluorescent Protein (EGFP) sequence in frame (Fig. 1b). Finally, we also targeted the start codon of the *Foxg1* gene to generate a conditional KI by inserting a Lox-Stop-Lox (LSL) cassette upstream of a triple FLAG sequence (Fig. 1c).

The size of the transgenes ranged from ∼ 700bp to 2.5kb, and 5 hours infection with high titers of recombinant AAV6 was performed for all KIs (a summary of the strategy used for the three KI approaches is detailed in Table 1).

**Table 1.**
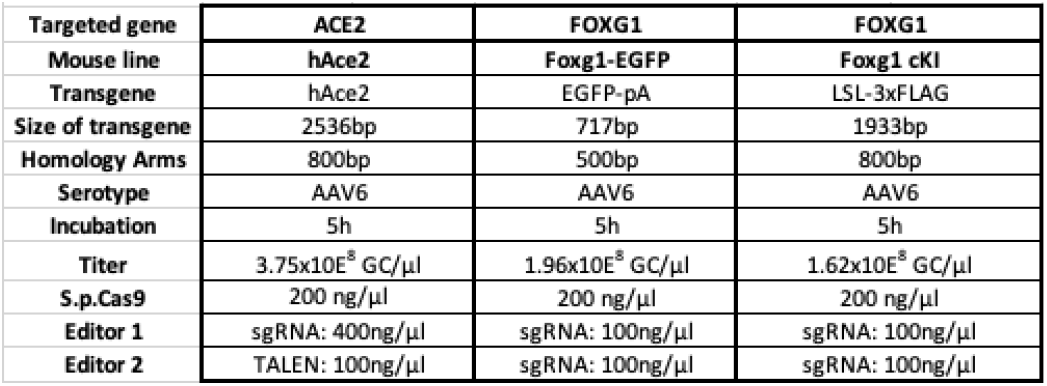
CRISPR-READI strategy to generate the KI mouse lines. Details of the transgenes and editing specifications used to generate the mouse lines by AAV-driven gene editing. bp = base pairs, GC/µl = genome-copy per microliter, ng/µl = nanogram per microliter.

### PCR genotyping of the KI mice does not identify any illegitimate mutation

#### hAce2 KI mice

To identify potential founders, we first ran a transgene-specific PCR. Out of eight pups, four (50%) carried the transgene (Fig. 2a). To validate the insertion of the transgene at the endogenous *Ace2* locus, we then performed 5’ and 3’ junction PCRs (Fig. 2b-c). Out of four potential founders, three carried the transgene at the insertion site. The fourth one (#43215) may display random insertion or episomal presence of the transgene. PCR genotyping did not reveal any apparent illegitimate event at the targeted site. We next selected two founders (#43204 and 43205) to establish the colonies.

**Figure 2.**
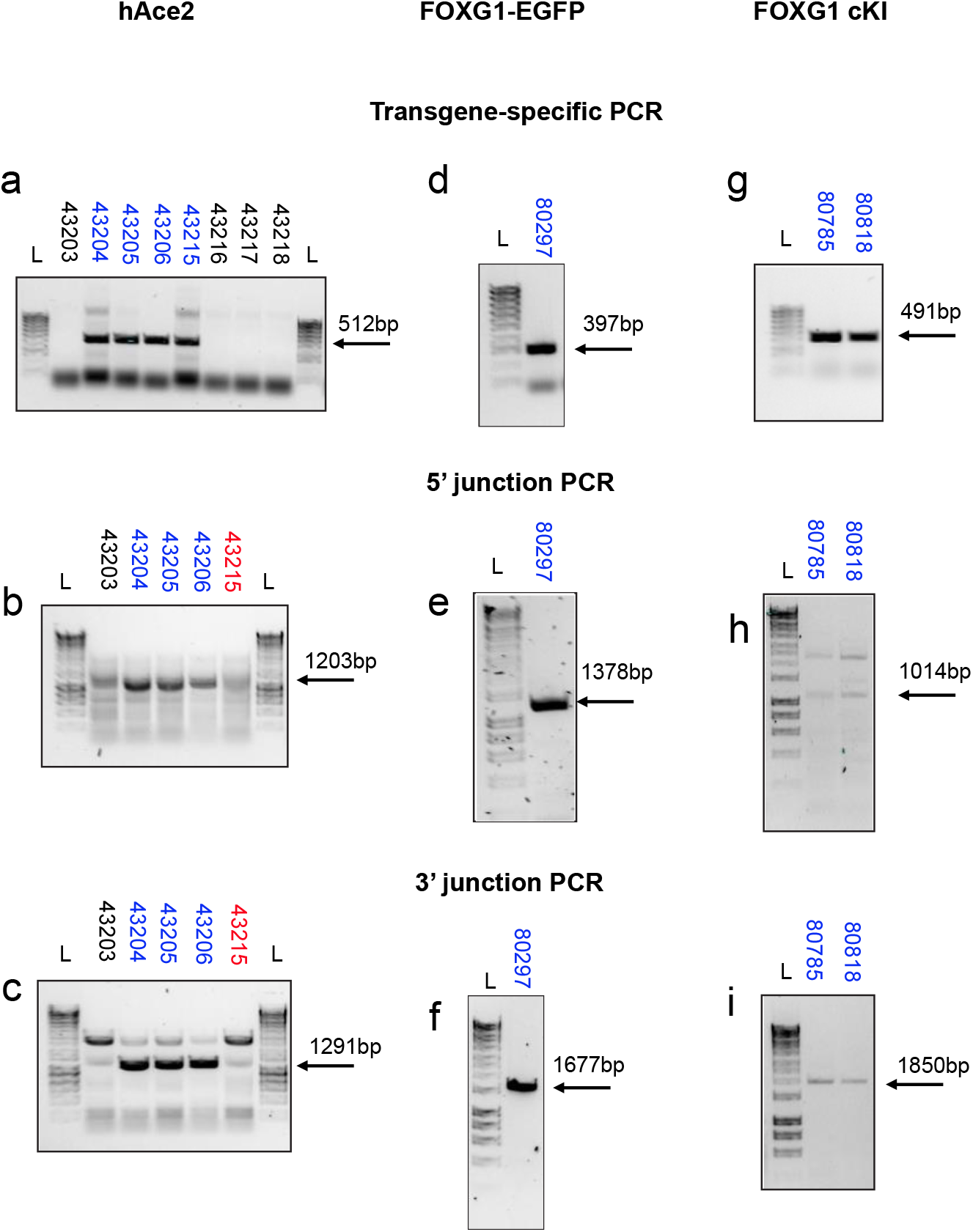
PCR Genotyping outcome for the KI mouse lines. Illustrative gel electrophoresis for the three KI mouse lines generated by CRISPR-READI. (a) Transgene-specific genotyping of the hAce2 founders (G0) showed that 4 out of 8 pups generated (50%) carried the hAce2 transgene. Targeted integration of the hAce2 transgene was confirmed by 5’ (b) and 3’ (c) junction PCRs. Founder #43215 may carry the AAV template as an episome. (d) Transgene-specific genotyping of the selected Foxg1-EGFP line. Targeted integration for the Foxg1-EGFP line was confirmed by 5’ (e) and 3’ (f) junction PCRs. (g) Transgene-specific genotyping of the selected Foxg1 conditional KI (Foxg1 cKI) line. Targeted integration for the Foxg1 cKI line was confirmed by 5’ (h) and 3’ (i) junction PCRs. L = ladder.

#### Foxg1 KI mice

Similar to the hAce2 KI mice, the AAV-driven gene editing was highly efficient at generating Foxg1 KI mice. Indeed, transgene-specific PCR identified sixteen potential founders out of 41 live pups (39%) for the Foxg1-EGFP; and three potential founders out of 18 live pups (17%) for the Foxg1 cKI, respectively (data not shown).

We selected one Foxg1-EGFP founder and two Foxg1 cKI founders and bred them with wildtype C57BL/6J mice to establish the colonies. We performed transgene-specific and junction PCRs on F1 mice and observed a normal genotyping profile with expected band sizes for all Foxg1 KI mice (Fig. 2 d-i). The sequence of all primers used in this study can be found in Supplemental Table 1.

**Supplementary table 1.**
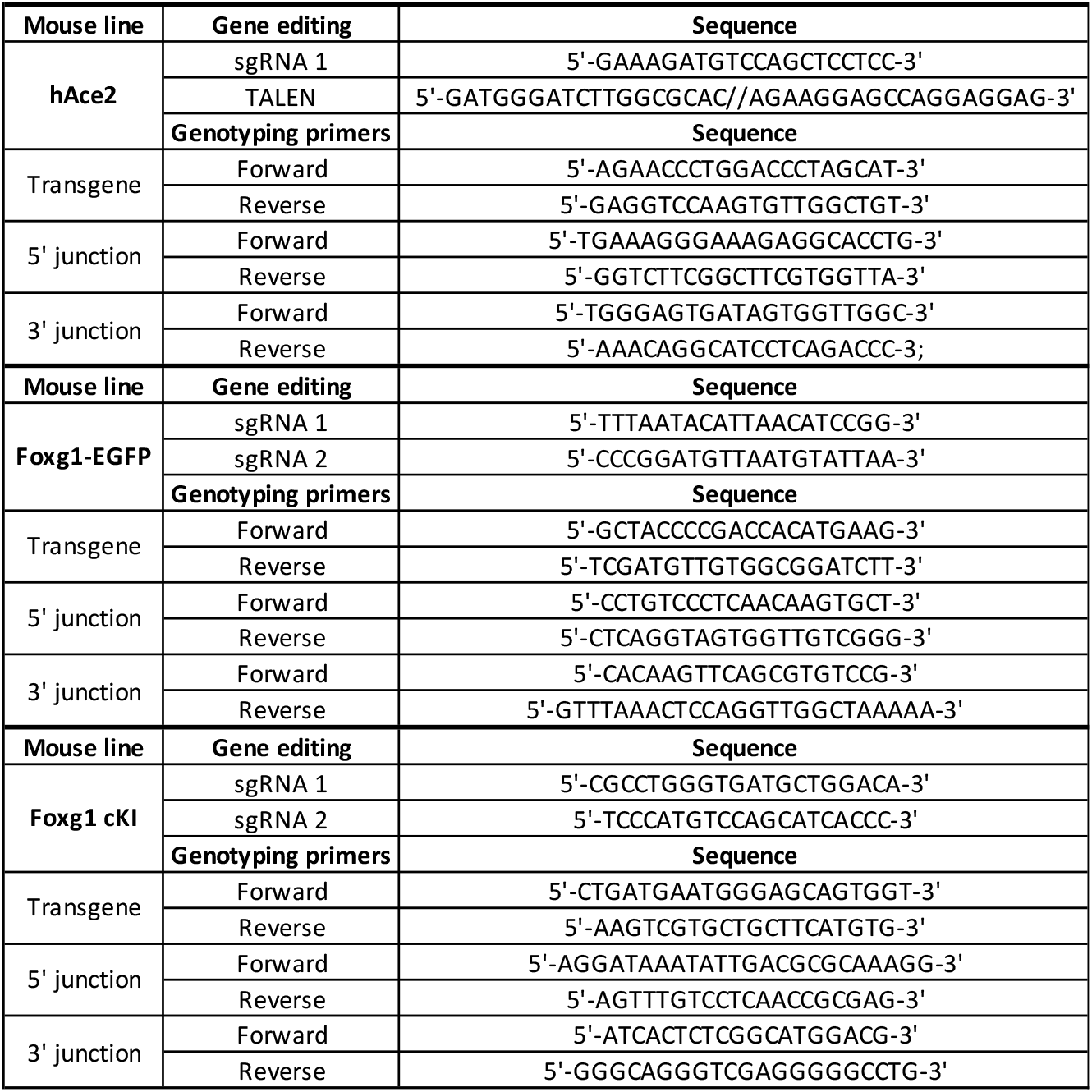
Sequences of the editing components and PCR primers used in this study.

### Cas9 enrichment generates adequate level of coverage

We performed genomic enrichment for five mice, two homozygous (#64005 and #65209) and three heterozygous (#80297, #88164 and #88312), and designed four sgRNAs for each locus (see *Methods*) to enrich the genomic regions of interest (ROI), following the nanopore Cas9-targeted sequencing (nCATS) method (Gilpatrick et al. 2020). The enrichment targeting a ∼3-5kb ROI centered around the integration site (Fig. 3a-c), yielded an on-target read depth ranging from 11 to 252x, representing 0.24-2.28% of all reads (Fig. 3a-c and Supplemental Table 2). Most samples yielded over 40x coverage, enough to reconstruct the KI consensus sequences using a *de novo* assembly approach, except for mouse #64005, which only achieved 11x coverage.

**Supplementary table 2.** Sequencing outputs following Cas9-enriched nanopore sequencing. Details of the on-target (i.e., containing the region of interest), off-target (i.e., not containing the region of interest) and total reads produced on the MinION for the five selected mice. N50 = sequence length of the shortest contig at 50% of the total assembly length.

| #Sample | Target | On-target |  |  | Off-target |  |  | All |  |  |
| --- | --- | --- | --- | --- | --- | --- | --- | --- | --- | --- |
|  |  | Total_reads | Total_base | N50 | Total_reads | Total_base | N50 | Total_reads | Total_base | N50 |
| 64005 | Ace2 | 11 | 128429 | 14005 | 4471 | 88654381 | 35125 | 4510 | 88836096 | 35068 |
| 65209 | Ace2 | 47 | 262843 | 5387 | 14313 | 291583183 | 36571 | 14425 | 291881756 | 36559 |
| 80297 | Foxg1-EGFP | 252 | 467781 | 2167 | 83140 | 933728501 | 21227 | 83726 | 934345287 | 21213 |
| 88164 | Foxg1 cKI | 221 | 1519782 | 9865 | 12383 | 162510756 | 37459 | 12672 | 164083342 | 37002 |
| 88312 | Foxg1 cKI | 99 | 488754 | 5506 | 4104 | 20167721 | 15515 | 4336 | 20772634 | 14286 |

**Figure 3.**
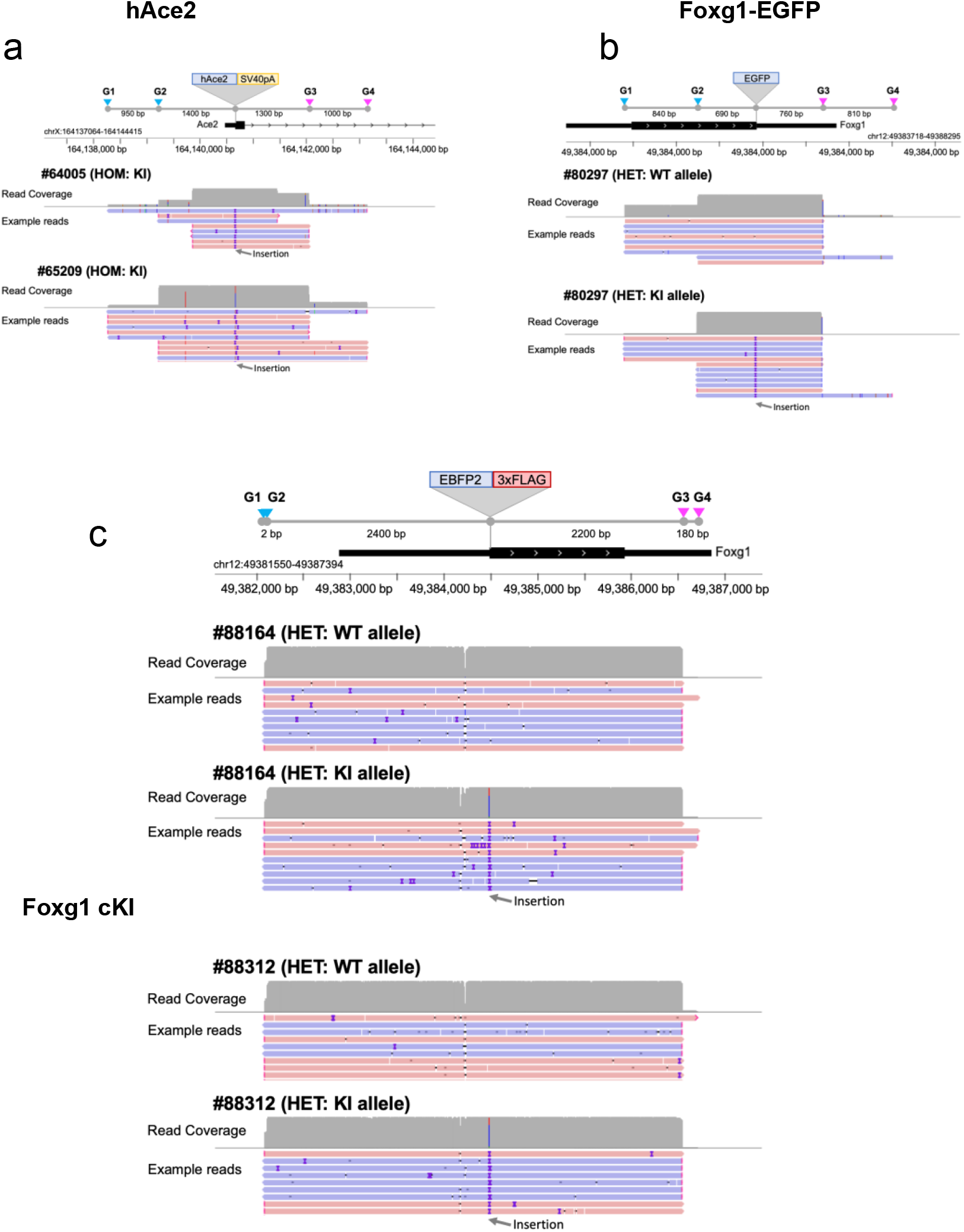
Cas9 enrichment strategy and resulting read depth for each KI. Illustration of the Cas9-based enrichment (nCATs method) of genomic DNA using four sgRNA and resulting read coverage. (a) Enrichment of the murine Ace2 targeted region using four sgRNA encompassing the insertion site of the hAce2 transgene. Example of reads for each homozygous hACe2 mouse tested (#64005 and #65209). (b) Enrichment of the murine Foxg1 targeted region using four sgRNA encompassing the insertion site of the EGFP transgene. Example of reads for the heterozygous Foxg1-EGFP mouse tested (#80297). (c) Enrichment of the murine Foxg1 targeted region using four sgRNA encompassing the insertion site of the EBFP2-3xFLAG transgene. Example of reads for the two heterozygous Foxg1cKI mouse tested (#88164 and #88312). G1-G4 = sgRNA1 to sgRNA4, WT = wildtype, KI = knock-in.

To evaluate whether adaptive sampling (AS) (Martin et al. 2022) could enhance coverage post Cas9-enrichment, two mice (#64005 and #65209) were sequenced, both with and without AS (as referenced in Table 2).

**Table 2.**
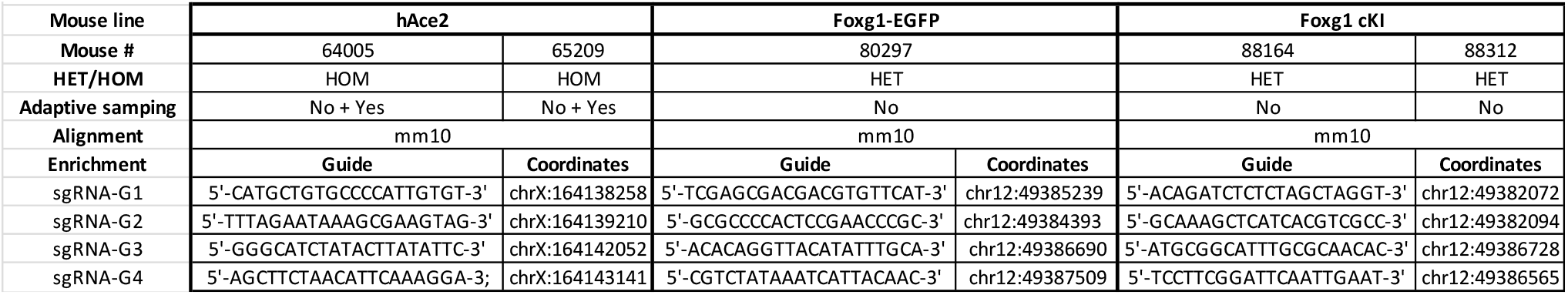
Cas9 enrichment strategy for nanopore long read sequencing. The nanopore Cas9-targeted sequencing (nCATS) method requires enrichment of the genomic sequence of interest. This table details the sequence of the guides used to enrich specific regions centered around the transgene integration site. HOM = homozygous, HET = heterozygous, chr = chromosome.

AS was found to increase the number of on-target reads in both cases, allowing the production of a valid consensus sequence for mouse #64005. However, the degree of improvement following AS was only moderate, with an increase in depth from 1.3 to 4.7 times (Table 3). Nonetheless, Cas9 enrichment alone yielded a sufficient number of reads (Supplemental Table 2) across most experiments for *de novo* assembly, presenting a viable option to generate consensus sequences in scenarios with limited computational resources (adaptive sampling requires a workstation equipped with a high-performance GPU).

**Table 3.** Effect of adaptive sampling on sequencing outputs following Cas9-enriched nanopore sequencing. Details of the on-target (i.e., containing the region of interest) and off-target (i.e., not containing the region of interest) reads produced on the MinION for the same mice with and without adaptive sampling (AS). N50 = sequence length of the shortest contig at 50% of the total assembly length.

| Table 3: effect of adaptive sampling on sequencing outputs following Cas9-enriched nanopore sequencing |  |  |  |  |  |  |  |  |
| --- | --- | --- | --- | --- | --- | --- | --- | --- |
| #Sample | Cas9 | Adaptive sampling | On-target |  |  | Off-target |  |  |
|  |  |  | Total_reads | Total_base | N50 | Total_reads | Total_base | N50 |
| 64005 | Yes | No | 11 | 128429 | 14005 | 4471 | 88654381 | 35125 |
| 64005 | Yes | Yes | 52 | 654184 | 14019 | 17966 | 10652182 | 494 |
| 65209 | Yes | No | 47 | 262843 | 5387 | 14313 | 291583183 | 36571 |
| 65209 | Yes | Yes | 61 | 303334 | 5378 | 19601 | 10280310 | 504 |

### Long read sequencing identifies occurrences of concatemerization

All nCATS libraries were sequenced on either a MinION Mk1c or a GridION. The analysis of the consensus sequences was performed by aligning them to the mm10 reference assembly. Dot plots visualization (Fig. 4) revealed the presence of concatemers at the insertion site.

**Figure 4.**
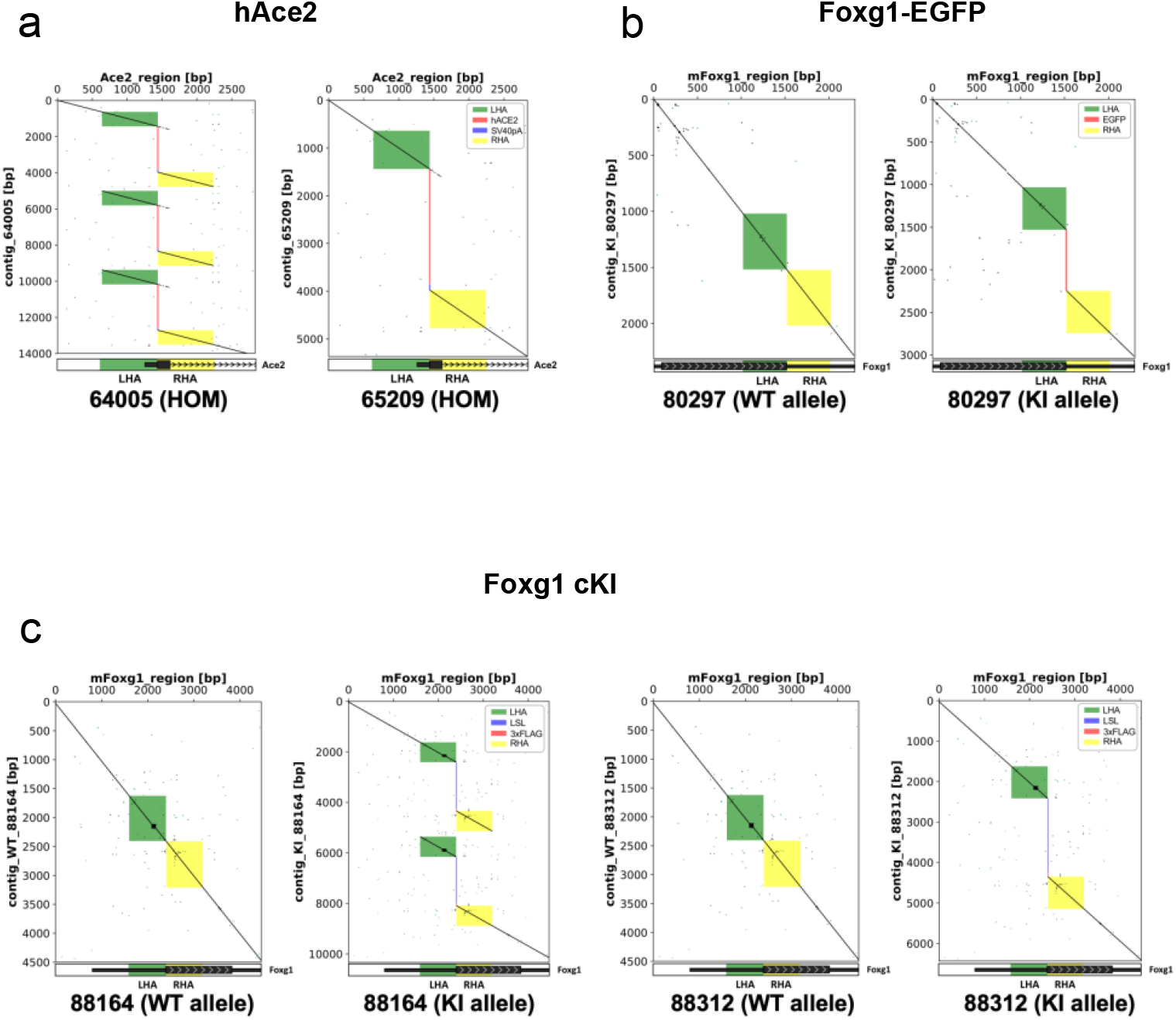
Visualization of the editing outcomes at the targeted loci following AAV-driven gene editing. Dot plots of the targeted loci shot that 2 lines out of 5 (40%) carry an array of multicopies (i.e., concatemers) at the insertion site. x axis = theoretical KI sequences, y axis = consensus sequences obtained by nanopore sequencing. A continuous line indicates full alignment, a broken line indicates a mismatch. (a) Dot plots analysis for the hAce2 homozygous mice reveals that mouse #65209 carries one copy whereas mouse #64005 carries three copies of the transgene. (b) Dot plots analysis for the Foxg1-EGFP heterozygous mouse reveals that mouse #80297 carries one copy of the transgene. (c) Dot plots analysis for theFoxg1 cKI heterozygous mice reveals that mouse #88164 carries two copies whereas mouse #88132 carries one copy of the transgene. LHA = left homology arm, RHA = right homology arm, ITR = inverted terminal repeat sequence, hAce2 = human Ace2 coding sequence, SV40pA = simian virus 40 polyadenylation signal, EGFP = Enhanced Green Fluorescence Protein sequence, 3xFLAG = triple FLAG tag, LSL = Lox-Stop-Lox.

Indeed, mouse #64005 (homozygous hAce2) carried three copies of the transgene, while mouse #88164 (Heterozygous Foxg1 cKI) had two copies integrated at the cut site. Overall, two mice out of five (40%) carried concatemers at the integration site (Table 4).

**Table 4.**
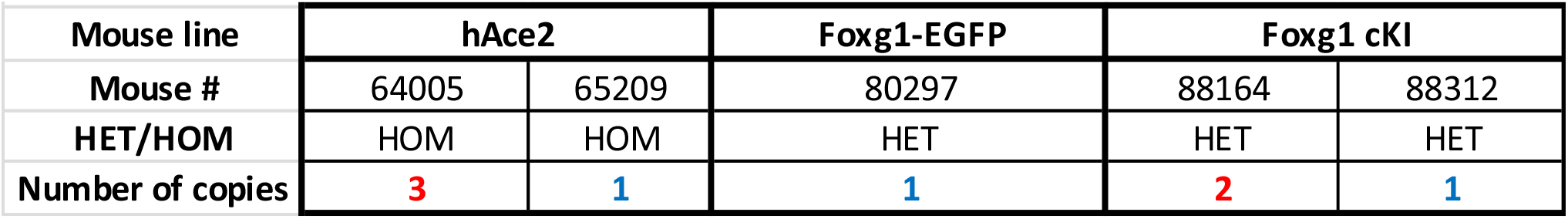
Long read sequencing outcome for the selected KI mice. Details of the number of copies integrated at the insertion site for the five selected mice. Note that two out of five (40%) mice carry multicopy (i.e., concatemers).

| Table 4: Long read sequencing outcome for the selected KI mice |  |  |  |  |  |
| --- | --- | --- | --- | --- | --- |
| Mouse line | hAce2 |  | Foxg1-EGFP | Foxg1 cKI |  |
| Mouse # | 64005 | 65209 | 80297 | 88164 | 88312 |
| HET/HOM | HOM | HOM | HET | HET | HET |
| Number of copies | 3 | 1 | 1 | 2 | 1 |

### Partial backbone integration at the insertion site

When aligning the consensus sequences of the selected mice to the sequence of the AAV plasmids used to generate the KIs, we did not find any backbone integration in the mice that carried single copies inserted. Conversely, both mice carrying concatemers also carried part of the backbone, essentially the Inverted Terminal Repeat (ITR) sequences (Fig. 5). Mouse #64005 contained both ITRs, whereas only the 3’ ITR was detected in the genome of mouse #88164. Importantly, these partial integrations were not found at the outermost extremities, where homologous directed repair occurred, and as such were not detectable in junction PCRs.

**Figure 5.**
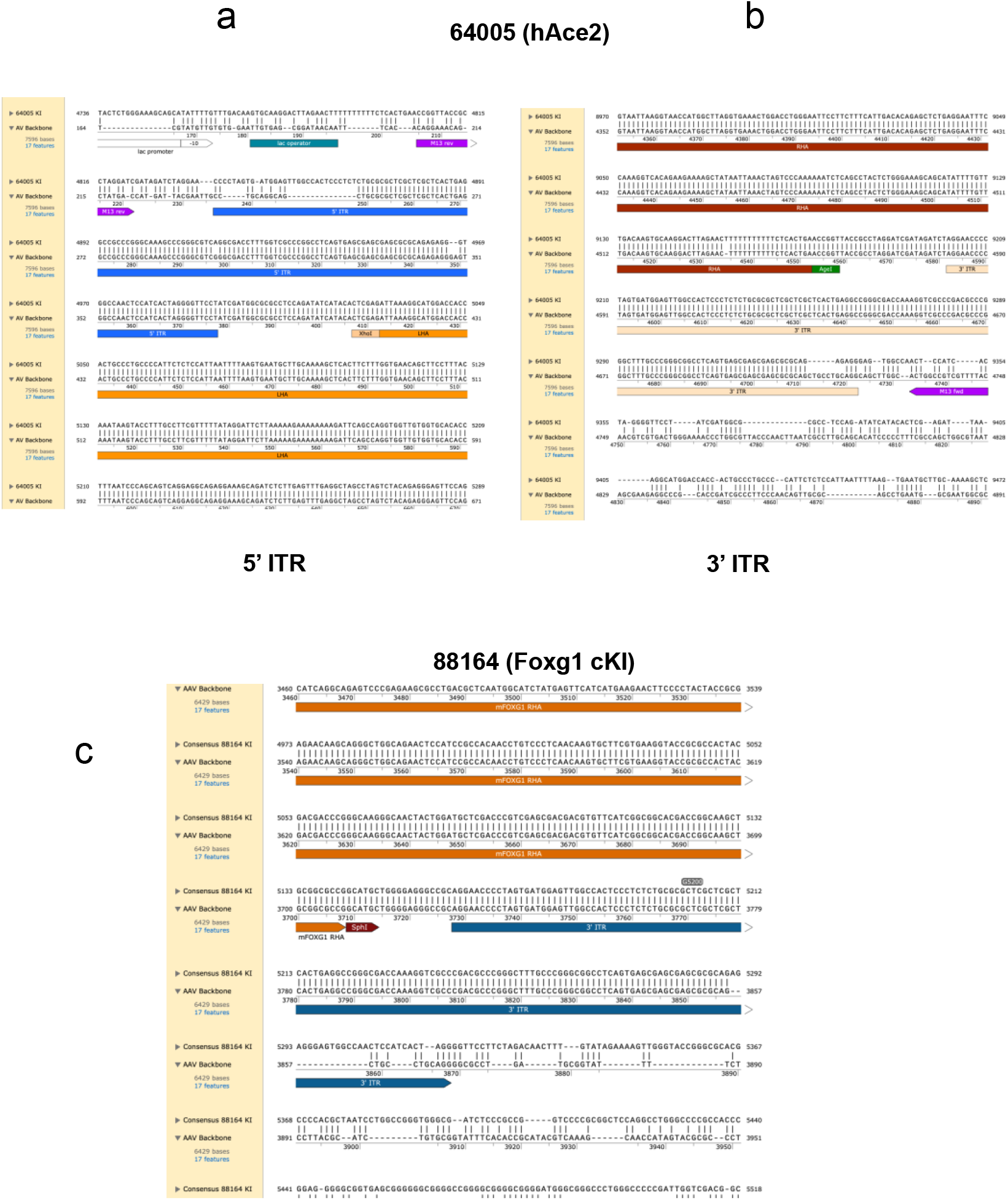
Alignment of the consensus sequences with the respective AAV plasmids. The alignment identifies partial insertion of backbones in concatemer carriers. The alignment between the hAce2 concatemer carrier consensus sequence (mouse #64005) and the sequence of the AAV plasmid used to generate this KI reveals that both the 5’ITR (a) and the 3’ ITR (b) sequences align partially, illustrating the integration of these ITR sequences together with a small part of the backbone containing the cloning sites (i.e., XhoI and AgeI). The alignment between the Foxg1 cKI concatemer carrier consensus sequence (mouse #88164) and the sequence of the AAV plasmid used to generate this KI reveals that the 3’ ITR sequence (c) aligns partially, illustrating the integration of this ITR sequence together with a small part of the backbone containing the cloning site (i.e., SphI). Note that the alignment of mice carrying single copies and wildtype alleles did not show any integration of the backbone. LHA = left homology arm, RHA = right homology arm, ITR = inverted terminal repeat sequence

## Discussion

Gene editing in mice to generate models of human disorders is critical to biomedical research. For instance, the insertion of the human Ace2 cDNA into its murine counterpart allows for the “humanization” of the *Ace2* gene, contributing to the development of new therapeutics against Sars-CoV-2 injections (Sun et al. 2020). Similarly, rare genetic disorders such as Foxg1 syndrome require the development of mouse models to study the pathomechanisms underlying the disorders (Younger et al.).

Although the disruption of genomic sequences (knock-out) is relatively straightforward in zygotes, the targeted insertion of large transgenes following homologous directed repair (HDR) remains challenging. Recently, AAV-driven electroporation of CRISPR RNP complexes in mouse zygotes proved a reliable and seamless method to generate KI mice, and transgenic cores around the world routinely use this method to generate mouse models (Duddy et al. 2024).

However, quality control (QC) for these models is typically done by transgene-specific and junction PCRs, followed by Sanger sequencing. Yet this method has been shown to be inadequate to detect potential on-target mutations (Simkin et al. 2022). Although off-target mutations are typically rare (Peterson et al. 2023) and no more frequent than genetic drift in mice (Iyer et al. 2015; Nakajima et al. 2016; Peterson et al. 2023), on-target mutations may occur with high frequency. To this end, we performed Long Read Sequencing (LRS) of the targeted loci following CRISPR-READI and found occurrences of on-target illegitimate mutations. Out of five mice analyzed by LRS, two carried multicopy integration (i.e., concatemers). Furthermore, the alignment of the consensus sequences with that of the respective AAV plasmids revealed partial integration of the AAV backbone, essentially the Inverted Terminal Repeat (ITR) sequences. Such illegitimate on-target mutations were typically not detected when we used the traditional PCR genotyping strategy (Mizuno et al. 2018), because these mutations did not occur at the outermost boundaries of the integration site. Instead, these sequences were typically found in between copies of the transgene, suggesting that the concatemerization occurred before integration.

When comparing the efficacy of Cas9 enrichment alone versus its combination with adaptive sampling, our data suggested that adaptive sampling could enhance coverage. Therefore, we advocate for the use of adaptive sampling when resources permit. Nonetheless, Cas9 enrichment alone yielded a sufficient number of reads across most experiments, presenting a viable option in scenarios with limited computational resources.

The entire process, from DNA extraction to data analysis, was completed within 4 days. This work highlights the potential of this method to provide a rapid and efficient means of assessing transgene insertion in a timely manner. Moreover, this approach has the potential to be widely applied as a tool for identifying and characterizing structural rearrangements and repetitive regions following gene editing in mice.

Although the present analysis is restricted to a limited number of mice, it has been previously demonstrated that genomic double-stranded breaks tend to capture foreign DNA (Sargent et al. 1997). As such, concatemers and/or illegitimate integrations have been observed with high frequency in different settings, including human cells edited with ZFN (Olsen et al. 2010; Radecke et al. 2010), murine embryonic stem cells edited with CRISPR (Erbs et al. 2023), *C. Elegans* (Dickinson et al. 2013) and Zebrafish (Gutierrez-Triana et al. 2018) edited with CRISPR, and cattle edited with TALEN (Norris et al. 2020). Even a typical workflow of microinjection of plasmids with CRISPR RNPs can generate up to 60% of the lines carrying multicopy integrations in mice (Skryabin et al. 2020). This could only be mitigated by biotinylation of the donor template (Medert et al. 2023). Importantly, high level of AAV vector integration (up to 47%) was also found when performing AAV-driven electroporation of CRISPR RNP complexes in cultured murine cells (Hanlon et al. 2019) and human hepatocytes (Ginn et al. 2020), in line with our findings in zygotes.

Therefore, we recommend using long read sequencing as a stringent QC for KI lines generated using CRISPR-READI, and potentially other methods. This work highlights the importance of in-depth validation of the mutant lines generated by transgenic cores, which is critical to ensure reproducibility of animal research, and as such, helps prevent animal wastage. Besides, this work also yields important information with regards to the uptake of this new method, given that CRISPR-Cas9/rAAV6 therapeutic strategies are currently implemented for monogenic diseases such as severe combined immunodeficiency (SCID) (Iancu et al. 2023).

## Methods

### Generation of KI mouse lines

The generation of KI mice was performed following the CRISPR-READI method (Chen et al. 2019). Briefly, C57BL/6J fertilized zygotes were infected for 5 hours with AAV6 carrying the respective transgene of interest in modified culture medium (Embryotech Laboratories, ETECH-EL) before thorough washing and ex-vivo electroporation of CRISPR reagents, as previously reported (Morey et al. 2022). Recombinant AAV6 was commercially sourced (Vector Builder, pilot scale packaging >2×10^11^ GC/ml). The zygotes were then reimplanted into pseudopregnant outbred mice (ARC(s), Ozgene) following conventional protocols (Delerue and Ittner 2017).

#### hAce2 KI mice

The first exon of the Ace2 gene (ENSEMBL ENSMUSG00000015405) was targeted using a commercially synthesised (Integrated DNA Technologies, Inc) guide (sequence in Supplemental Table 1). This single-guide RNA (sgRNA) was rationally designed using a computational tool (Oliveros et al. 2016) to minimize off-targets (https://bioinfogp.cnb.csic.es/tools/breakingcas/) and incubated for 10min at room temperature with *Alt-R™ S.p. Cas9* Nuclease *V3* (IDT # 1081058) to form ribonucleoprotein (RNP) complexes. TALEN mRNA (ThermoFisher Scientific) targeting the same locus was also added to the editing mix.

Following AAV infection, this editing mix was electroporated (NEPA21, Nepagene) into fertilised C57BL/6 zygotes with the respective concentrations: 200ng/μl Cas9, 400ng/μl sgRNA, 100ng/μl TALEN (Table 1).

#### Foxg1-EGFP KI mice

The first exon of the Foxg1 gene (ENSEMBL ENSMUSG00000020950) was targeted at the stop codon using two commercially synthesised (Integrated DNA Technologies, Inc) guides (sequences in Supplemental Table 1). These single-guide RNAs (sgRNAs) were rationally designed using a computational tool (Oliveros et al. 2016) to minimize off-targets (https://bioinfogp.cnb.csic.es/tools/breakingcas/) and incubated for 10min at room temperature with *Alt-R™ S.p. Cas9* Nuclease *V3* (IDT # 1081058) to form ribonucleoprotein (RNP) complexes. Following AAV infection, this RNP mix was electroporated (NEPA21, Nepagene) into fertilised C57BL/6 zygotes with the respective concentrations: 200ng/μl Cas9, 100ng/μl each guide (Table 1).

#### Foxg1 cKI mice

These mice were generated using the exact same procedure (and concentrations) as the one used for the Foxg1-EGFP mice, however the sgRNAs targeted the start codon of the first exon of the Foxg1 gene.

For all KI lines, live pups were produced and bred to establish colonies. PCR genotyping on isopropanol-precipitated DNA from tail biopsies was performed to identify potential founders. Genotyping screen consisted in transgene-specific PCR, followed by 5’ and 3’ junctions PCRs (primer sequences in Supplemental Table 1).

### Genomic DNA extraction

High Molecular Weight (HMW) DNA was obtained using the Monarch^®^ HMW DNA extraction kit for Tissue (T3060L, New England Biolabs), as previously described (Low et al. 2022). Briefly, fresh kidney tissues from heterozygous (HET) or Homozygous (HOM) mice were harvested, kept on ice and 20mg of tissue was immediately processed.

### Library preparation

Enrichment of the genomic region of interest (ROI) was performed using the Cas9 Sequencing Kit (SQK-CS9109, ONT^©^) following the nCATS method (Gilpatrick et al. 2020). Briefly, following dephosphorylation, Cas9-guided adapter ligation and dA tailing were conducted on 5μg of HMW genomic DNA for each mouse, using four rationally designed sgRNAs (Labun et al. 2016) (two upstream and two downstream of the ROI, approximately 0.7 to 2.4 kilobases from the insertion site (Fig. 3a-c). Next, AMPure XP bead purification was performed, and final concentrations were measured using Qubit fluorometric quantification (Koetsier and Cantor 2021) (Q33238, ThermoFisher Scientific) before loading each nCATS library on an R9.4.1 flow cell (FLO-MIN106D, ONT^©^).

### Long Read Sequencing

Where applicable, adaptive sampling was employed to selectively capture reads belonging to the target regions of interest (ROI) defined in the corresponding BED files (see *Supplemental Material*) specifying 15 kilobases upstream and downstream from the ROI, using the *Mus musculus* C57BL/6J (mm10) as genomic reference. Nanopore sequencing was performed with or without adaptive sampling for 72h on a MinION Mk1b or GridION without reloading. Fast5 files and FASTQ files were collected for analysis.

### Data Analysis and visualization

Base calling was executed using GUPPY (v6.4.6) with the High-accuracy Model (HAC). The resulting FASTQ files were then aligned to the mouse reference genome (mm10) using minimap2 (v2.24). The depth of reads targeting specific regions was assessed using BEDTools genomecov (v2.30). The reads that mapped to the target region encompassing sgRNAs cleavage sites were extracted using the intersect function of BEDTools (v2.30). These reads were then used for *de novo* assembly with Canu (v2.2) followed by subsequent polishing with nanopore FASTQ using two rounds of Flye-polishing (v2.9.2-b1786) and one round of Medaka-polishing (v1.8.0). Visualization of on-target reads and assembly outcomes was performed using the Integrative Genomics Viewer (IGV v2.16.2). Finally, dot plot graphs were generated using FlexiDot (v1.06) to analyse the editing outcomes at the insertion sites, while consensus sequences were aligned to the AAV plasmid sequences using Snapgene v7.1.1. (Fig. 5).

## Data access

All data that support the findings of this study are available upon request. The raw sequence reads generated in this study have been submitted to the NCBI BioProject database under accession number PRJNA1076264 (http://www.ncbi.nlm.nih.gov/bioproject/1076264).

## Acknowledgements

This work was supported by a joint research seeding grant from Macquarie University and Mahidol University (round 2022). Further research support was from the Australian Research Council (DP210101957). Pattaraporn Nimsamer received financial support from the NSRF via the Program Management Unit for Human Resources & Institutional Development, Research Innovation (Grant No. B13F660073). The computing facilities were supported by Mahidol University and the Office of the Ministry of Higher Education, Science, Research and Innovation under the Reinventing University project : the Center of Excellence in AI-Based Medical Diagnosis (AI-MD) sub-project, NSTDA Supercomputer center (ThaiSC), and the National e-Science Infrastructure Consortium.

## Author contributions

F.D. and T.W. designed the study; M.W.L., P.J., N.L., P.N. and F.D. performed the experiments, M.W.L., P.J., J.S., N.L., M.C., T.W. and F.D. analysed data; T.W. and F.D. supervised the study; L.M.I., T.W. and F.D. obtained funding; F.D. wrote the first draft of the manuscript. All authors edited the manuscript.

## Ethics statement

All procedures were approved by Macquarie University Animal Ethics Committees and were conducted in accordance with the Australian Code of Practice for the Care and Use of Animals for Scientific Purposes.

